# *ZMPSTE24* Missense Mutations that Cause Progeroid Diseases Decrease Prelamin A Cleavage Activity and/or Protein Stability

**DOI:** 10.1101/198069

**Authors:** Eric D. Spear, Ehr-Ting Hsu, Laiyin Nie, Elisabeth P. Carpenter, Christine A. Hrycyna, Susan Michaelis

**Affiliations:** Department of Cell Biology, The Johns Hopkins School of Medicine, Baltimore, MD 21205; Department of Chemistry, Purdue University, West Lafayette, IN 47907; Structural Genomics Consortium, University of Oxford, Oxford, UK

## Abstract

The human zinc metalloprotease ZMPSTE24 is an integral membrane protein critical for the final step in the biogenesis of the nuclear scaffold protein lamin A, encoded by *LMNA*. After farnesylation and carboxyl methylation of its C-terminal CAAX motif, the lamin A precursor, prelamin A, undergoes proteolytic removal of its modified C-terminal 15 amino acids by ZMPSTE24. Mutations in *LMNA* or *ZMPSTE24* that impede this prelamin A cleavage step cause the premature aging disease Hutchinson-Gilford Progeria Syndrome (HGPS) and the related progeroid disorders mandibuloacral dysplasia-type B (MAD-B) and restrictive dermopathy (RD). Here we report a “humanized yeast” system to assay ZMPSTE24-dependent cleavage of prelamin A and examine the eight known disease-associated *ZMPSTE24* missense mutations. All show diminished prelamin A processing and fall into three classes, with defects in activity, protein stability, or both. Notably, some ZMPSTE24 mutants can be rescued by deleting the E3 ubiquitin ligase Doa10, involved in ER-associated degradation of misfolded membrane proteins, or by treatment with the proteasome inhibitor bortezomib, which may have important therapeutic implications for some patients. We also show that ZMPSTE24-mediated prelamin A cleavage can be uncoupled from the recently discovered role of ZMPSTE24 in clearance of ER membrane translocon-clogged substrates. Together with the crystal structure of ZMPSTE24, this “humanized yeast system” can guide structure-function studies to uncover mechanisms of prelamin A cleavage, translocon unclogging, and membrane protein folding and stability.

## INTRODUCTION

The integral membrane <u>z</u>inc <u>m</u>etallo<u>p</u>rotease ZMPSTE24 plays a critical role in human health and longevity through its role in the maturation of the nuclear scaffold protein lamin A from its precursor, prelamin A (Bergo et al., 2002; Michaelis and Hrycyna, 2013; Pendas et al., 2002). Mature lamin A, together with nuclear lamins B and C, contributes to the structural integrity and proper functioning of the nucleus (Butin-Israeli et al., 2012; Dechat et al., 2010; Dittmer and Misteli, 2011; Dorado and Andres, 2017; Gerace and Huber, 2012; Gruenbaum and Foisner, 2015; Mattout et al., 2006). Defects in prelamin A processing by ZMPSTE24 are a primary cause of progeria (Capell and Collins, 2006; Davies et al., 2009; Gordon et al., 2014; Michaelis and Hrycyna, 2013). The premature aging disorder Hutchinson-Gilford Progeria syndrome (HGPS; OMIM #176670) results from mutations in the *LMNA* gene (encoding prelamin A) that block ZMPSTE24 processing, while the related progeroid diseases mandibuloacral dysplasia-type B (MAD-B; OMIM #608612) and restrictive dermopathy (RD; OMIM #275210) result from *ZMPSTE24* mutations that diminish protease function (Davies et al., 2009; De Sandre-Giovannoli et al., 2003; Eriksson et al., 2003; Navarro et al., 2014). Understanding the mechanistic details of prelamin A processing by ZMPSTE24 is thus critical for designing therapeutic approaches for these progeroid diseases and may also provide insights into the normal physiological aging process.

The posttranslational maturation of prelamin A is a multi-step process. Prelamin A contains a C-terminal CAAX motif (C is cysteine, A is usually an aliphatic amino acid, and X is any residue). Like other CAAX proteins, prelamin A undergoes a series of three reactions, referred to as CAAX processing (Fig. 1, Steps 1-3) which includes farnesylation of cysteine, proteolytic removal of the - AAX residues mediated redundantly by ZMPSTE24 or RCE1, and carboxyl methylation of the farnesylated cysteine by isoprenylcysteine methyltransferase (ICMT) (Davies et al., 2009; Michaelis and Barrowman, 2012; Michaelis and Hrycyna, 2013; Wang and Casey, 2016). Prelamin A is distinct from all other CAAX proteins in higher eukaryotes in that following CAAX processing, prelamin A undergoes a second endoproteolytic cleavage event uniquely mediated by ZMPSTE24 (Fig. 1, Step 4). This second cleavage removes the C-terminal 15 amino acids, including the modified cysteine, to yield mature lamin A (Bergo et al., 2002; Pendas et al., 2002). In progeroid disorders, this second ZMPSTE24-promoted cleavage of prelamin A is compromised, leading to the accumulation of a permanently farnesylated and carboxyl methylated form of prelamin A, which is the toxic “culprit” in disease (Davies et al., 2009; Gordon et al., 2014; Worman et al., 2009).

**Figure 1.**
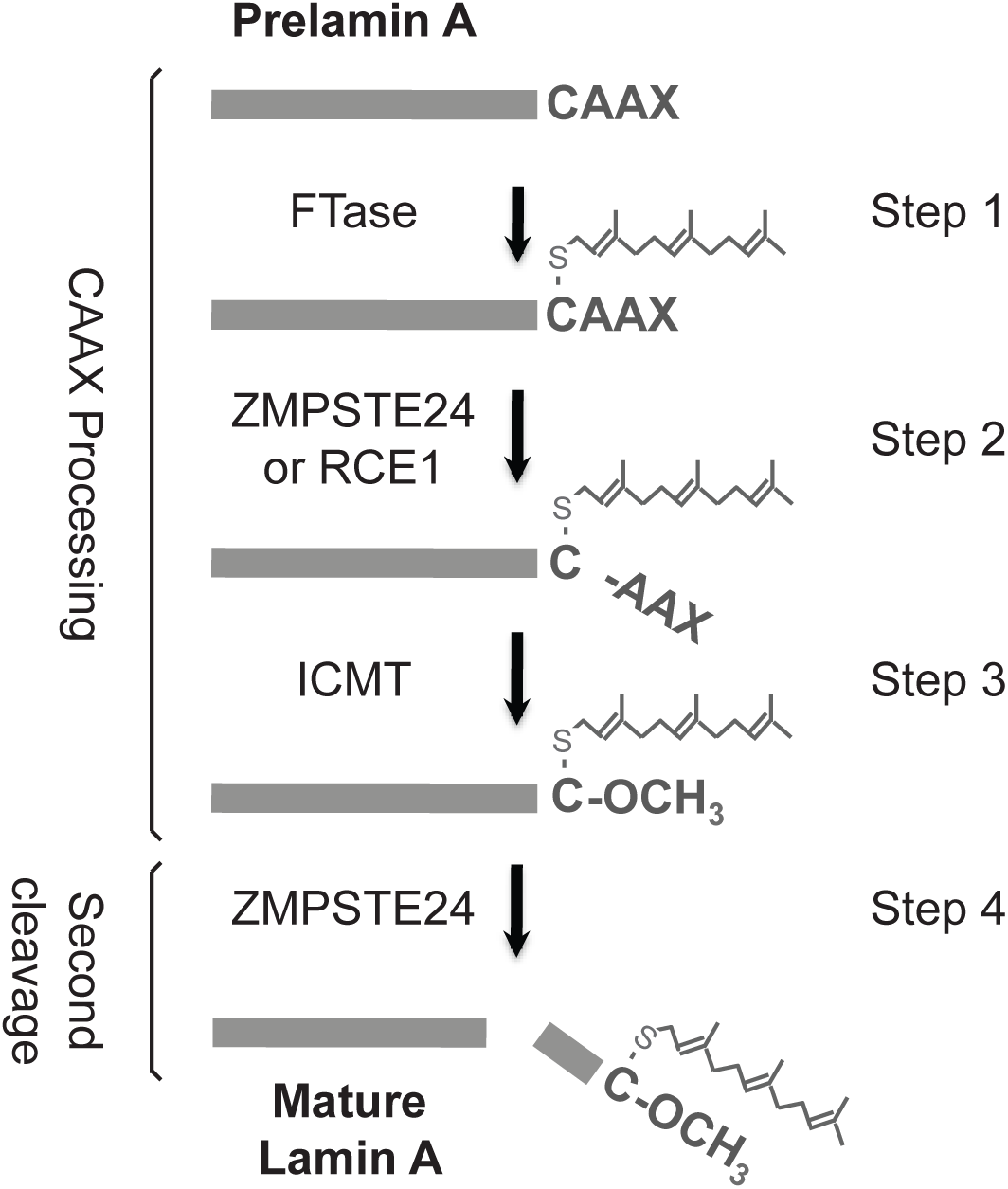
The prelamin A biogenesis pathway. The four steps of prelamin A posttranslational processing shown here are described in the text. The lipid farnesyl (a 15-carbon-long isoprenoid lipid) and the carboxyl methyl group (O-CH_3_) are indicated. The enzymes that mediate CAAX processing are shown: farnesyltransferase (FTase), the proteases ZMPSTE24 and Ras converting enzyme (RCE1), and the isoprenylcysteine carboxylmethyl transferase (ICMT). It should be noted that while Step 2 in CAAX processing can be carried out redundantly for prelamin A either by ZMPSTE24 or RCE1, Step 4 of prelamin A processing is solely mediated by ZMPSTE24. When ZMPSTE24 is absent, processing is blocked at Step 4 and not Step 2, since RCE1 is present ((Varela et al., 2008) and Hrycyna, C. Hsu, E.T., and Michaelis, S, unpublished data).

MAD-B, HGPS, and RD represent a spectrum of disorders of increasing severity (Barrowman et al., 2012b; Navarro et al., 2014). In HGPS, the best studied of these, children manifest accelerated aging symptoms starting at one year of age, including failure to thrive, lipodystrophy, hair loss, joint ailments, and cardiovascular disease, and they typically die in their mid-teens from heart attack or stroke. Nearly all HGPS patients harbor a dominant *LMNA* mutation that, through altered splicing, generates an internally deleted version of prelamin A called progerin, which retains its CAAX motif but lacks the ZMPSTE24 cleavage site, and causes disease phenotypes (De Sandre-Giovannoli et al., 2003; Eriksson et al., 2003; Gordon et al., 2014; Merideth et al., 2008). RD and MAD-B are due to recessive mutations in *ZMPSTE24*, and result from the accumulation of full-length prelamin A that is permanently farnesylated and carboxyl methylated. RD is far more severe than HGPS, being fatal at or before birth, and is due to complete loss of ZMPSTE24 function resulting from null mutations (frameshifts, premature termination, or large deletions) in both copies of *ZMPSTE24* (Moulson et al., 2005; Navarro et al., 2005; Navarro et al., 2014; Smigiel et al., 2010). In contrast, MAD-B is generally milder than HGPS, with patients having variable survival rates and disease severity, yet all exhibiting lipodystrophy as a major disease phenotype. Individuals with MAD-B have one *ZMPSTE24* null allele and one *ZMPSTE24* missense allele that provides reduced but residual function (Table 1) (Agarwal et al., 2003; Barrowman et al., 2012b; Navarro et al., 2014). In general, the severity of these three progeroid diseases reflects the amount of permanently farnesylated and carboxyl methylated prelamin A that accumulates per cell. Recently individuals with metabolic syndrome and nonalchoholic fatty liver disease (NAFLD), both lipodystrophy-associated disorders, were also found to have a *ZMPSTE24* missense mutation (Table 1) (Brady et al., 2017; Dutour et al., 2011; Galant et al., 2016). In addition, diminished ZMPSTE24 processing of prelamin A may be important in normal aging, based on a study showing that prelamin A accumulation occurs in blood vessels from aging, and not young, individuals (Ragnauth et al., 2010). Because of the importance of diminished ZMPSTE24 processing of prelamin A in progeroid disease and possibly during normal aging, understanding the detailed mechanism of ZMPSTE24 is an important area of research.

**Table 1.** ZMPSTE24 missense mutations that result in MAD-B or atypical progeria

| **Partial**<br>***zmpste24***<br>**mutation** | **Partial/Null**<br>***zmpste24***<br>**mutation** | **Disease** | **Accumulates**<br>**prelamin A** | **ZMPSTE24**<br>**levels** | **References** |
| --- | --- | --- | --- | --- | --- |
| L94P | L94P | MAD-B | Yes | Decreased | (Ben Yaou et al., 2011) |
| P248L | Q41X | MAD-B | Yes | ND | (Akinci et al., 2017)<br>(Miyoshi et al., 2008) |
| P248L | W450X | MAD-B | Yes | ND | (Ahmad et al., 2010)<br>(Akinci et al., 2017) |
| N265S | Y70S(fsX3) | MAD-B | ND | ND | (Cunningham et al., 2010) |
| N265S | L362F(fsX18) | AT “HGPS” | Yes | ND | (Shackleton et al., 2005) |
| N265S | L362F(fsX18) | MAD-B | ND | ND | (Agarwal et al., 2006) |
| W340R | L362F(fsX18) | MAD-B | ND | ND | (Agarwal et al., 2003)<br>(Moulson et al., 2005) |
| Y399C | Y399C *(LOH) | MAD-B | ND | ND | (Haye et al., 2016) |
| L425P | L425P | MAD-B | Yes | ND | (Harhour et al., 2016) |
| L438F | none detected | Metabolic syndrome | Yes | Normal | (Dutour et al., 2011)<br>(Galant et al., 2016) |
| L438F | none detected | Nonalcoholic fatty liver disease (NAFLD) | ND | ND | (Brady et al., 2017) |
| L462R | none detected | RD | ND | ND | (Thill et al., 2008) |

\*LOH is loss of heterozygosity

ZMPSTE24 is widely conserved in eukaryotes ranging from yeast to mammals (Barrowman and Michaelis, 2009; Pryor et al., 2013; Quigley et al., 2013). The *Saccharomyces cerevisiae* homolog Ste24 is the founding member of this family and was discovered based on its role in the proteolytic maturation of the secreted yeast mating pheromone a-factor (Barrowman and Michaelis, 2011; Boyartchuk and Rine, 1998; Fujimura-Kamada et al., 1997; Michaelis and Barrowman, 2012; Tam et al., 1998). Prelamin A and the a-factor precursor are distinct from other CAAX proteins, as they are the only ones that undergo additional cleavage by ZMPSTE24/Ste24 after CAAX processing is completed (Barrowman and Michaelis, 2011; Beck et al., 1990; Bergo et al., 2002; Pendas et al., 2002; Sinensky et al., 1994). ZMPSTE24 and its homologs contain seven transmembrane spans and a consensus zinc metalloprotease HEXXH motif (H is histidine, E is glutamate, X is any amino acid) which is critical for coordinating zinc and performing catalysis (Barrowman et al., 2012b; Quigley et al., 2013). The recently solved X-ray crystallography structure of human ZMPSTE24, and that of the virtually superimposable yeast Ste24, reveal it to be a completely novel class of protease (Clark et al., 2017; Pryor et al., 2013; Quigley et al., 2013). The seven helical spans of ZMPSTE24 form a voluminous intramembrane “hollow” chamber, with the HEXXH catalytic domain positioned such that it faces the interior of the chamber, with a side portal(s) in the chamber presumably providing a site for prelamin A entry. This unusual ZMPSTE24 structure raises important functional questions, including how ZMPSTE24 mediates specificity for prelamin A access into its chamber, what residues are involved in positioning prelamin A for its cleavage(s), and what might be the role of ZMPSTE24’s large chamber. The answers to these questions are of fundamental biological interest and will help us to understand how specific ZMPSTE24 disease alleles malfunction and might be corrected. They may also shed light on how certain HIV protease inhibitors such as lopinavir are able to inhibit ZMPSTE24 (Coffinier et al., 2007; Mehmood et al., 2016). Such insights could also have relevance to physiological aging.

ZMPSTE24 is dually localized in the inner nuclear and ER membranes (Barrowman et al., 2008) and performs cellular functions in addition to its well-established role in the proteolytic maturation of prelamin A and a-factor. Recent work shows that ZMPSTE24 plays a protein quality control role by clearing “clogged” Sec61 translocons of post-translationally secreted proteins that have aberrantly folded while in the process of translocation (Ast et al., 2016; Kayatekin et al., 2018). Intriguingly, a role for ZMPSTE24 in defending cells against a wide variety of enveloped viruses, independent of its catalytic activity, has also been recently reported (Fu et al., 2017).

Because of ZMPSTE24’s importance in human health and disease and its novel structure, it would be advantageous have a high throughput system to probe structure-function relationships in this protease. Here we report a “humanized yeast system” to specifically assay the second ZMPSTE24 cleavage step in prelamin A maturation (Fig. 1, Step 4). We show that the eight currently known disease-causing ZMPSTE24 missense alleles (Table 1) all have decreased prelamin A cleavage *in vivo* and fall into distinct classes-those that affect solely cleavage activity, those that affect *in vivo* protein stability through ER-associated degradation (ERAD) by the ubiquitin-proteasome system (UPS), and those that affect both. Notably, for two unstable ZMPSTE24 disease mutants, P248L and W340R, when ubiquitylation or proteasome activity is blocked, both their stability and catalytic activity are significantly restored. These findings have implications for therapeutic strategies that could ultimately optimize “personalized medicine” approaches. The *in vivo* assay system we present here, along with the ease of gene manipulation and genetic strategies available in yeast, hold promise for future high-throughput structure-function studies on ZMPSTE24.

## RESULTS

### ZMPSTE24 can perform the upstream cleavage of its *bona fide* substrate prelamin A in yeast

We previously showed that human ZMPSTE24 could functionally replace its yeast homolog Ste24 for the proteolytic maturation of its non-native substrate, the yeast mating pheromone a-factor (Barrowman and Michaelis, 2009; Schmidt et al., 2000; Tam et al., 1998). We also developed an assay in which the extent of yeast mating broadly correlated with the severity of ZMPSTE24 disease alleles, such that those that cause RD (null alleles) show more severe mating defects than those that cause the milder disease MAD-B (missense alleles) (Barrowman et al., 2012b). However, the mating assay is less than ideal for the mechanistic dissection of ZMPSTE24 because it relies on ZMPSTE24-dependent cleavage of the cross-species substrate a-factor, and because it cannot distinguish between ZMPSTE24’s two cleavage activities. Because the unique step in prelamin A cleavage by ZMPSTE24 is the second cleavage, and because it is the lack of this step that causes progeroid diseases, we set out to develop a system to specifically measure this ZMPSTE24-mediated processing step for its *bona fide* substrate prelamin A, which is not normally present in S. *cerevisiae*.

To create a ‘humanized’ yeast system to study ZMPSTE24-dependent processing of prelamin A, we expressed a C-terminal segment from the human prelamin A protein (amino acids 431-664) (Fig. 2A), which contains all the necessary signals for CAAX processing and the ZMPSTE24-dependent unique cleavage (Barrowman et al., 2008; Barrowman et al., 2012a). To serve as size markers for comparison, we also constructed a mutant prelamin A, L647R, that is known to be uncleavable by ZMPSTE24 in mammalian cells (Barrowman et al., 2012a; Hennekes and Nigg, 1994; Mallampalli et al., 2005; Wang et al., 2016) as well as a version expressing the correctly processed mature form of prelamin A (amino acids 431-646). All versions were N-terminally tagged with 10His-3myc to allow detection by western blotting and were integrated into the yeast genome. In this humanized yeast system ZMPSTE24 is expressed from a low-copy number yeast CEN plasmid and is N-terminally 10His-3HA tagged (Fig. 2A). Prelamin A cleavage is measured by quantitation of the mature and prelamin A species present in cells at steady state. Importantly, our strain background retains Rce1 to allow efficient –AAXing (Fig. 1, Step2), thus eliminating any effect that mutant ZMPSTE24 proteins may have on this first cleavage step.

**Figure 2.**
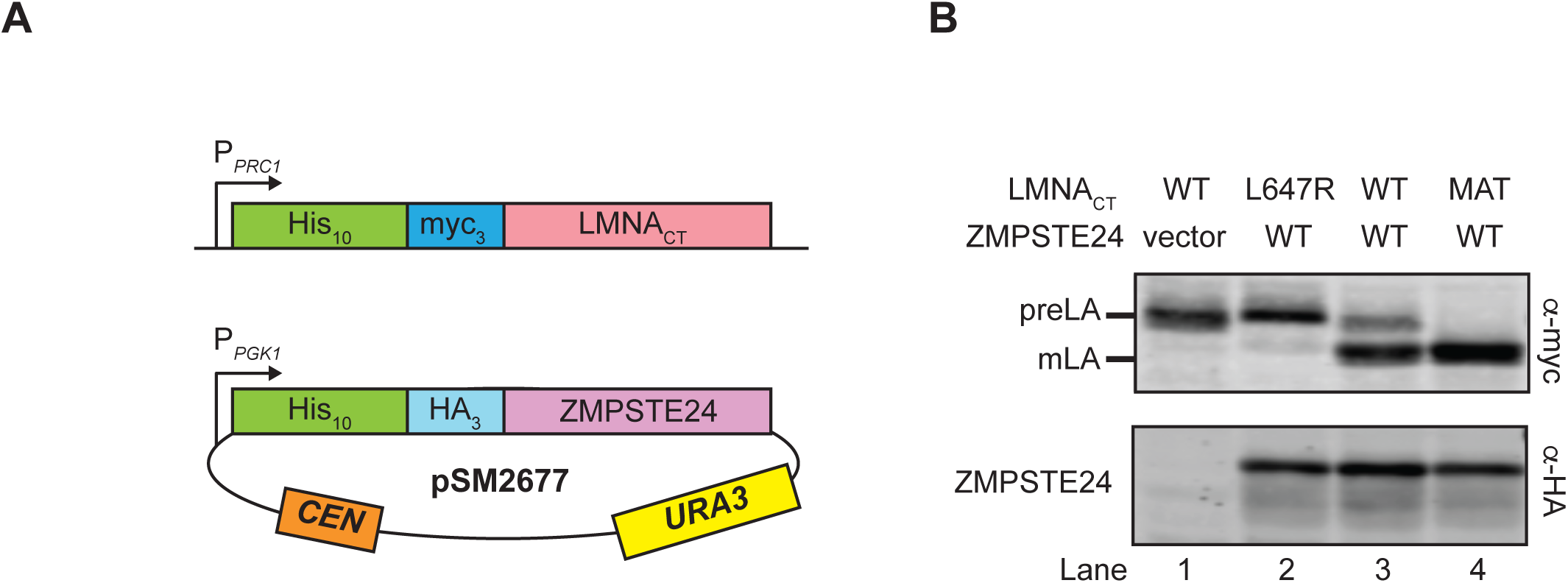
Prelamin A is processed to mature lamin A by human ZMPSTE24 in yeast. (A) A schematic of the humanized yeast system is shown. The prelamin A model substrate contains amino acids 431-664 from the C-terminus of human *LMNA* (referred to LMNA_CT_) fused to a 10His-3myc epitope tag. It is expressed from the *PRC1* promoter (*P_PRC1_*) and is chromosomally integrated into a *ste24*Δ strain background, resulting in strain SM6158. Full-length human ZMPSTE24 with an N-terminal 10His-3HA epitope tag is expressed from the *PGK1* promoter *(P_PGK1_)* on a *CEN URA3* plasmid (pSM2677; (Barrowman et al., 2012b)). (B) Lysates from *ste24*Δ strains expressing wild-type prelamin A (Lanes 1 and 3), uncleavable prelamin A (Lane 2, L647R) or mature lamin A (Lane 4, MAT) and human ZMPSTE24 (Lanes 2, 3 and 4) or vector alone (Lane 1) were analyzed for prelamin A processing by SDS-PAGE and western blotting. Prelamin A (preLA) and mature lamin A (mLA) were detected with anti-myc antibodies; ZMPSTE24 was detected with anti-HA antibodies. Strains in lanes 1-4 are SM6158/pRS316, SM6177/pSM2677, SM6158/pSM2677, and SM6178/pSM2677, respectively.

We first tested whether human ZMPSTE24 could process prelamin A in a *ste24*Δ strain. Plasmid-borne ZMPSTE24, but not vector alone, resulted in two bands observed by western blotting with anti-myc antibodies (Fig. 2B top panel, compare lanes 1 and 3). Importantly, these bands co-migrated with the “uncleavable” and “mature” forms (Fig. 2B top panel, lanes 2 and 4, respectively), suggesting that the prelamin A substrate was properly cleaved by ZMPSTE24. We note that yeast Ste24 can also cleave prelamin A (Fig. S1) and that prelamin A processing to the mature form by ZMPSTE24 can be enhanced by the addition of a second copy of ZMPSTE24 (Fig. S2).

We also tested whether prelamin A cleavage in yeast required the CAAX modifications farnesylation and carboxyl methylation, as it does in mammalian cells (Barrowman et al., 2008; Barrowman et al., 2012a). Wild-type ZMPSTE24, but not a catalytic-dead mutant, H335A, resulted in mostly mature lamin A (Fig. 3A, compare lanes 1 and 2). Mutation of the CAAX motif cysteine to a serine (C661S), which prevents its farnesylation, completely blocked ZMPSTE24-dependent cleavage of prelamin A (Fig. 3A, lane 3). The unmodified C661S prelamin A migrated slightly more slowly than farnesylated prelamin A in the H335A ZMPSTE24 mutant (Fig. 3A, compare lanes 2 and 3), as has been previously observed (Yang et al., 2008). We also examined prelamin A cleavage in a *ste14*Δ strain, which lacks the yeast ICMT. As observed in mammalian cells (Barrowman et al., 2008; Barrowman et al., 2012a; Young et al., 2005), blocking carboxyl methylation of the prelamin A substrate has a modest, but discernable effect on prelamin A cleavage (Fig. 3B, compare lane 3 to lane 1). Taken together, these experiments demonstrate the processing of prelamin A in yeast follows the same rules as in mammalian cells. Thus we can use our “humanized” yeast system to study ZMPSTE24-dependent processing of its *bona fide* substrate prelamin A.

**Figure 3.**
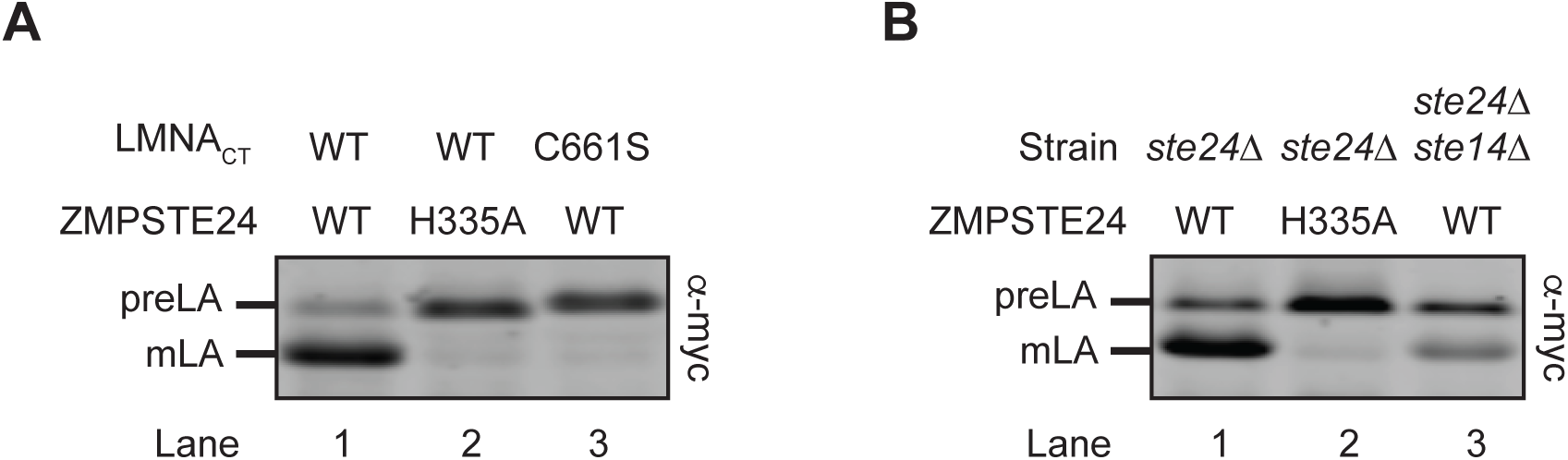
Cleavage of prelamin A in yeast, as in mammalian cells, requires farnesylation of the CAAX motif and is diminished when carboxyl methylation is lacking. (A) Prelamin A processing is blocked when farnesylation is absent. Prelamin A processing in *ste24*Δ strains expressing the indicated *LMNA*_CT_ (WT or C661S) and *ZMPSTE24* (WT and H335A) alleles was analyzed by SDS-PAGE and western blotting, as in Fig. 2. (B) The efficiency of prelamin A processing is reduced in a *ste14*Δ mutant strain. Prelamin A processing in strains expressing the indicated *ZMPSTE24* alleles was analyzed. Strains are *ste24*Δ only (Lanes 1 and 2) or a *ste24*Δ*ste14*Δ double mutant (Lane 3). Strains in lanes 1-3 are SM6158/pSM2677, SM6158/pSM2673, and SM6187/pSM2677, respectively.

### All ZMPSTE24 disease missense mutations show reduced prelamin A cleavage and some exhibit a low level of protein

One goal of developing a yeast *in vivo* cleavage assay was to determine whether particular *ZMPSTE24* disease alleles resulted in defective prelamin A cleavage, and by what mechanism(s), which ultimately might suggest therapeutic possibilities. Currently, eight different ZMPSTE24 substitution mutations are known to cause progeroid disorders (Table 1). We examined the processing efficiency of these alleles compared to WT ZMPSTE24. Also included in our panel are two mutations, H335A and H339A, known to abolish ZMPSTE24 activity by disrupting the zinc metalloprotease domain (H_335_EXXH_339_) (Barrowman and Michaelis, 2009; Barrowman et al., 2012b; Fujimura-Kamada et al., 1997; Quigley et al., 2013). As evident in Fig. 4A and summarized in Table 2, all of the mutations we examined showed reduced *in vivo* prelamin A cleavage compared to wild-type ZMPSTE24, albeit to widely varying degrees. For instance, L438F shows the highest residual activity at 57.2% of WT ZMPSTE24, while L462R shows the least at 6.5% (Fig. 4A, compare lanes 11 and 12 to lane 2). Notably, none of the disease alleles are as severe as the two catalytic dead mutants H335A and H339A (Fig. 4A, lanes 6 and 7), which have <2% WT ZMPSTE24 activity. Some of the mutations in our panel were previously shown to accumulate prelamin A in patient cells (L94P, P248L, N265S, L425P, and L438F; (Ben Yaou et al., 2011; Dutour et al., 2011; Harhouri et al., 2016; Miyoshi et al., 2008; Shackleton et al., 2005)), but directly comparing the extent of severity for these *ZMPSTE24* alleles was not possible in non-isogenic patient cells. The yeast system, however, is ideal for this purpose. Three of the mutants, W340R, Y399C, and L462R, had never been examined for prelamin A processing defects. Thus, our yeast system has for the first time confirmed a potential molecular basis underlying these latter mutants (prelamin A accumulation) and allows us to compare levels of residual processing between all known ZMPSTE24 disease alleles.

**Table 2.** ZMPSTE24 *in vivo* relative cleavage activity, steady state protein levels, and adjusted ZMPSTE24 enzyme activity (normalized to protein amount)

| **ZMPSTE24** | **Activity (% of WT)** | **Protein levels (% of WT)** | **Adjusted ZMPSTE24 activity (%)<sup>\*</sup>** | **Mutant Class<sup>\*\*</sup>** |
| --- | --- | --- | --- | --- |
| **vector** | **1.2** | **0** | **NA** | **-** |
| **WT** | **100** | **100** | **100** | **N/A** |
| **L94P** | **11.3** | **34.1** | **33.1** | **Class III** |
| **P248L** | **20.5** | **13.8** | **148.6** | **Class II** |
| **N265S** | **26.2** | **84.3** | **31.1** | **Class I** |
| **H335A** | **1.5** | **70.0** | **2.1** | **N/A** |
| **H339A** | **1.9** | **72.0** | **2.6** | **N/A** |
| **W340R** | **41.7** | **39.6** | **105.3** | **Class II** |
| **Y399C** | **25.8** | **92.9** | **27.8** | **Class I** |
| **L425P** | **41.9** | **62.1** | **67.5** | **Class III** |
| **L438F** | **57.2** | **67.9** | **84.2** | **Class III** |
| **L462R** | **6.5** | **29.7** | **21.9** | **Class III** |

<sup>\*</sup> Adjusted ZMPSTE24 activity = Activity/Protein levels X 100
<sup>\*\*</sup> Class I disease mutants are affected mainly in enzymatic function, Class II mainly in protein stability, Class III I both
<sup>\*\*\*</sup> Catalytic dead mutants are in red

**Figure 4.**
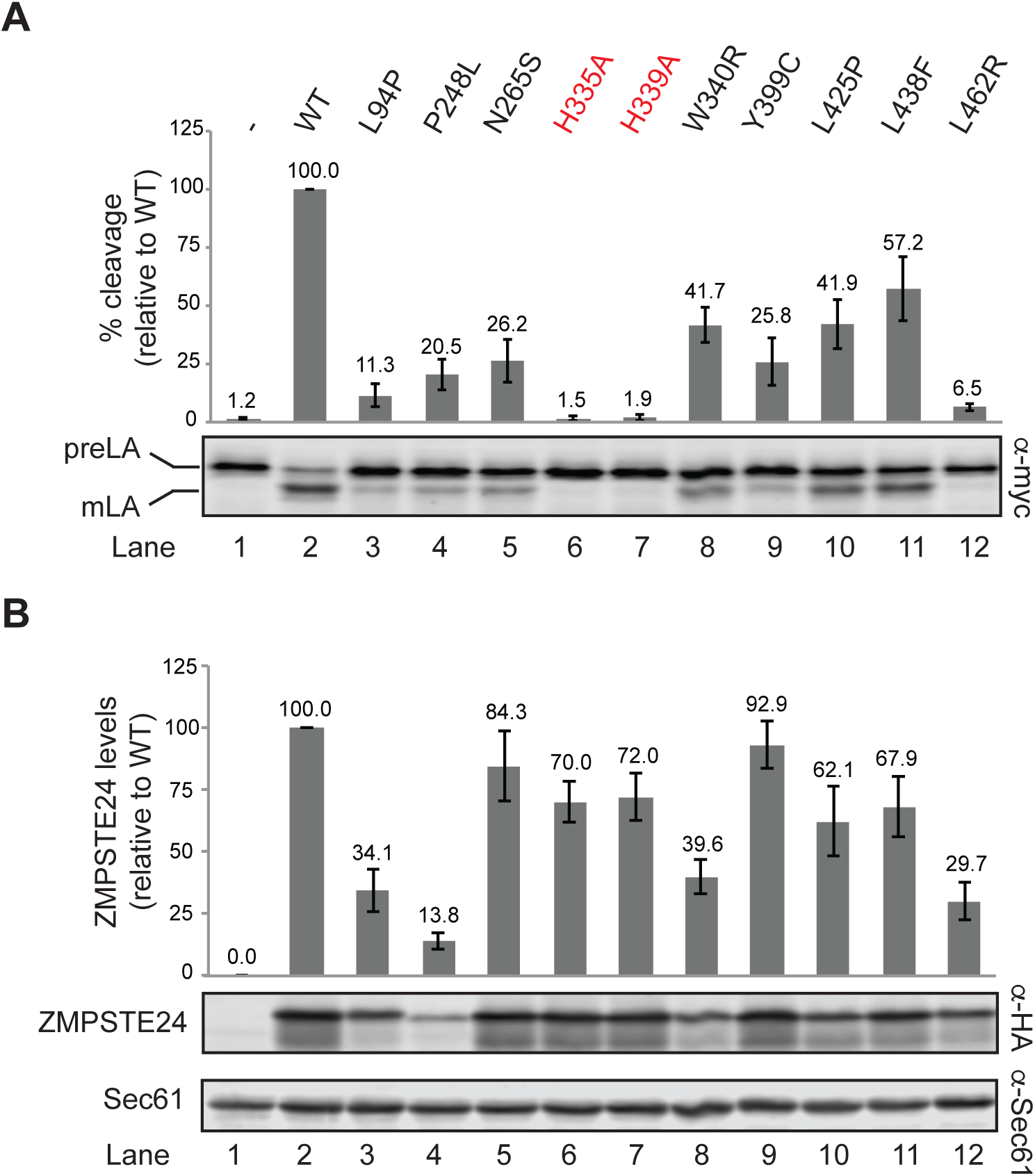
ZMPSTE24 disease mutants show diminished prelamin A cleavage, and for some alleles dramatically decreased protein levels. Lysates from strain SM6158 (*ste24*Δ *myc-LMNA*_CT_) transformed with plasmids expressing the indicated HA-ZMPSTE24 alleles or vector only were analyzed by SDS-PAGE and western blotting. (A) Average (mean) percentage of prelamin A cleavage for each ZMPSTE24 variant was calculated from 4 independent experiments, with standard deviation of the mean shown as error bars. For comparison, WT ZMPSTE24 cleavage was set to 100%. p<0.005 for all mutants compared to WT. (B) ZMPSTE24 proteins were detected with a-HA antibodies, and the ZMPSTE24 levels were normalized to the loading control Sec61, with WT ZMPSTE24 set to 100%. The average (mean) and standard deviation of the mean are shown for the same 4 experiments as in (A). p<0.05 for all mutants compared to WT, except N265S and Y399C, which were not considered to be significantly different from WT. We note that the multiple banding pattern seen here for ZMPSTE24 occurs not only in our yeast system, but also for endogenous ZMPSTE24 in mammalian cells (Pendas et al., 2002), and when ZMPSTE24 is expressed in other heterologous expression systems (Clark et al., 2017 and EP Carpenter and L. Nie, unpublished observations). The different mobilities may simply reflect distinct SDS-binding patterns for this protein or an as yet unknown modification.

Given the non-conservative amino acid substitutions in several of the ZMPSTE24 mutants, we considered the possibility that decreased prelamin A cleavage could at least in part be the result of ZMPSTE24 misfolding and subsequent degradation (Fig. 4B). Indeed, western blotting revealed varying amounts of ZMPSTE24 for some mutants. It should be noted that ZMPSTE24 resolves as a major band with a faster migrating smearing pattern and a minor band below it, as observed previously (39). Such anomalous SDS-PAGE migration patterns are not uncommon for membrane proteins and reflect the unusual detergent binding properties of their helical spans (Newman et al., 1981; Rath et al., 2009). Four of the mutants (L94P, P248L, W340R and L462R) showed steady-state ZMPSTE24 levels significantly less (<40%) than that of wild-type ZMPSTE24 (Fig. 4B, compare lanes 3, 4, 8 and 12 to lane 2). Notably, when prelamin A processing is normalized for the amount of ZMPSTE24 protein, two of the mutants, P248L and W340R, had an “adjusted ZMPSTE24 activity” of 100% or higher (Table 2), suggesting that degradation, and not compromised catalytic activity, are the problem for these alleles (and in the section below, we find these two mutants are significantly active when their degradation is blocked). Strikingly, however, other mutants, including N265S and Y399C, displayed near-normal ZMPSTE24 protein levels, yet retained only ~25-30% activity (Fig. 4A and B), which albeit less than WT, is far more than that of the catalytic dead mutants H335A and H339A (<2%) (Table 2).

Together, these results demonstrate that the humanized yeast system can differentiate three classes of ZMPSTE24 disease mutations: Class I mutants are those that affect mainly enzymatic activity (N265S and Y399C), Class II are those that affect mainly protein stability (P248L, W340R), and Class III are those that appear to affect both (L94P, L425P, L438F, and L462R).

### Blocking ubiquitylation and degradation of some ZMPSTE24 disease mutants rescues the prelamin A cleavage defect

Mutations in transmembrane proteins like ZMPSTE24 can result in degradation by the UPS, causing disease despite the fact that their catalytic function remains intact. In some cases, enhancing the folding or blocking the degradation of these mutant proteins using pharmacological chaperones, proteasome inhibitors, or mutants defective in ubiquitylation, can rescue protein levels enough to restore function within the cell (Amaral, 2015; Brodsky, 2012; Guerriero and Brodsky, 2012). We therefore asked whether blocking the ubiquitylation and degradation of the most unstable ZMPSTE24 mutant proteins, L94P, P248L, W340R and L462R, could rescue prelamin A cleavage.

To test whether blocking ubiquitylation affected ZMPSTE24 protein levels and activity, we deleted the gene encoding Doa10, a dually-localized ER and inner nuclear membrane E3 ligase known to ubiquitylate many misfolded transmembrane proteins and targets them for degradation (Deng and Hochstrasser, 2006; Huyer et al., 2004; Ravid et al., 2006; Swanson et al., 2001). Steady-state levels of the ZMPSTE24 mutant proteins increase ~2-5 fold in the *doa10*Δ strain, indicating all are substrates, either partially or fully, of this ubiquitin ligase (Fig. 5A, compare adjacent lanes for each disease mutant). Importantly, we observed that stabilization of P248L and W340R in the *doa10*Δ strain, but not L94P or L462R, restored prelamin A cleavage activity to near wild-type levels (Fig. 5B, compare adjacent lanes 5 and 6, 7 and 8, versus 3 and 4, 9 and 10).

**Figure 5.**
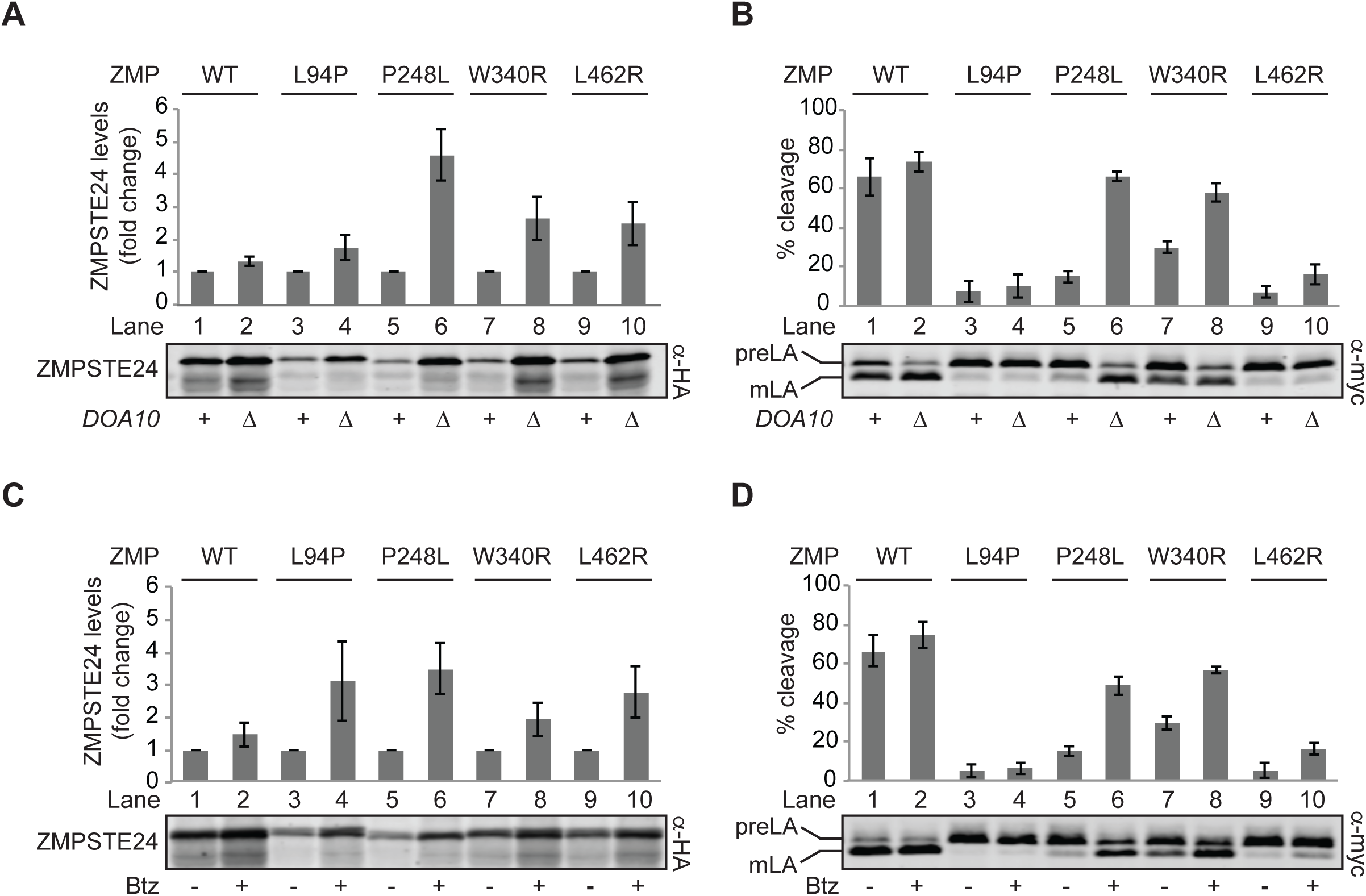
Blocking the ubiquitin/proteasome-dependent degradation of mutant ZMPSTE24 proteins enhances prelamin A cleavage for some ZMPSTE24 disease variants. The effects of blocking ubiquitylation are examined in (A and B), and proteasome inhibition in (C and D). Strains SM6158 (*ste24*Δ *myc-LMNA*_CT_) or SM6184 (*ste24*Δ*doa10*Δ *myc-LMNA*_CT_) expressing the indicated ZMPSTE24 variants were analyzed by SDS-PAGE and western blotting using α-HA (A) and a-myc (B) antibodies. ZMPSTE24 protein levels were normalized against the loading control Sec61 (not shown). The *doa10*Δ mutant strain is designated as “Δ” and the wild-type *DOA10* strain as “+”. Data shown is mean ± s.d. for four independent experiments. p<0.05 for all comparisons between + and Δ for ZMPSTE24 protein levels; p<0.005 for P248L and W340R comparing activity (B). To test the effect of proteasome inhibition on ZMPSTE24 protein levels and activity, strain SM6159 (*pdr5*Δ*ste24*Δ *myc-LMN*Δ_CT_) expressing the indicated ZMPSTE24 variants were treated with 20 µM bortezomib (+), or DMSO vehicle (−) as described in Materials and Methods. (C) HA-ZMPSTE24 proteins detected with anti-HA antibodies were normalized to the loading control Sec61 (not shown) and levels were expressed as fold change between treated (+) and untreated (−) samples. A representative gel is shown, with the mean ± s.d. for three independent experiments shown above. p<0.05 for all comparisons of mutant ZMPSTE24 proteins. (D) Prelamin A cleavage from the same samples shown in panel (C) was assessed with anti-myc antibodies. p<0.05 for P248L, W340R and L462R compared to WT.

We also tested whether the proteasome inhibitor bortezomib had similar effects as the *doa10*Δ mutant. Treatment of cells with 20 µM bortezomib for four hours resulted in ~2-4 fold more protein for all ZMPSTE24 mutants compared to drug vehicle alone (Fig. 5C, compare − and + lanes for each variant). In agreement with the above results where ubiquitylation is blocked, both P248L and W340R showed enhanced prelamin A cleavage upon proteasome inhibition (Fig. 5D, compare lanes 5 and 6, and lanes 7 and 8). Notably, these same two mutants, P248L and W340R, had shown WT or better “adjusted ZMPSTE24 activity” above (Table 2), when activity was normalized to the amount of ZMPSTE24 protein present. Taken together, these data suggest that some ZMPSTE24 patient mutations (Class II) are prematurely targeted for ubiquitin-mediated degradation, despite retaining catalytic activity. For patients with these alleles, therapeutic strategies that reverse the destruction of these otherwise functional enzymes by blocking their degradation could be beneficial.

The prelamin A cleavage and Sec61 translocon clearance functions of ZMPSTE24 can be genetically separated

Recently, yeast Ste24 and mammalian ZMPSTE24 were shown to have a specialized protein quality role for handling postranslationally secreted proteins that prematurely fold while translocating across the ER membrane, thereby “clogging” the Sec61 translocation machinery (Ast et al., 2016). Clearance of a reporter “clogger” protein by yeast Ste24 or heterologously expressed human ZMPSTE24 in *ste24*Δ yeast cells required their catalytic activity. Although four ZMPSTE24 disease alleles, including L94P, P248L, W340R and L438F, were previously assayed for their ability to clear the clogger substrate and shown to be defective to varying extents (Ast et al., 2016), four additional mutants in our current study were not tested.

The “clogger” reporter is a chimeric protein composed of the yeast glycoprotein Pdi1 fused to the clogging element bacterial dihydrofolate reductase (DHFR), followed by a stretch of N-linked glycosylation sequences (Ast et al., 2016). Its complete translocation into the ER lumen is observed as an SDS-PAGE mobility shift when all N-glycosylation sites (in both Pdi1 and the C-terminus) are modified. The hemi-glycosylated substrate is assumed to be partially translocated (clogged) and the unmodified protein represents a cytoplasmic pool that accumulates upon translocon clogging (Fig. 6). As observed previously, *ste24*Δ cells transformed with vector alone or the catalytic-dead ZMPSTE24 variants H335A and H339A had the most severe defects (Fig. 6, lanes 1, 6 and 7, respectively), with 36-44% of the reporter accumulating in the “clogged/cytoplasmic” forms, compared to only ~18% for wild-type ZMPSTE24 (Fig. 6, lane 2). Likewise, L94P and P248L, which have severe prelamin A processing defects also showed significant “clogged/cytoplasmic” accumulation (~30%) (Fig. 6, lanes 3 and 4). Surprisingly, however, other ZMPSTE24 mutants, including Y399C, L425P, and L438F, despite showing prelamin A cleavage defects, had little to no defects in “clogger” clearance (Fig. 6, lanes 5 and 9-11). L462R is also intriguing as it is relatively unstable and has a strong prelamin A cleavage defect, yet shows only a minor defect in “clogger” clearance (Fig. 6, lane 12). These findings suggest that while catalytic activity is required for both of ZMPSTE24’s functions, prelamin A processing and “declogging” may differ in important ways mechanistically. For instance, mutant ZMPSTE24 proteins may be differently affected in their ability to be recruited to the Sec61 translocon, or in their ability to permit access of the two different types of substrates (prelamin A vs. clogged proteins) into their active site chamber. It will be of interest to attempt to isolate ZMPSTE24 variants that can efficiently process prelamin A, but are defective for clogger clearance.

**Figure 6.**
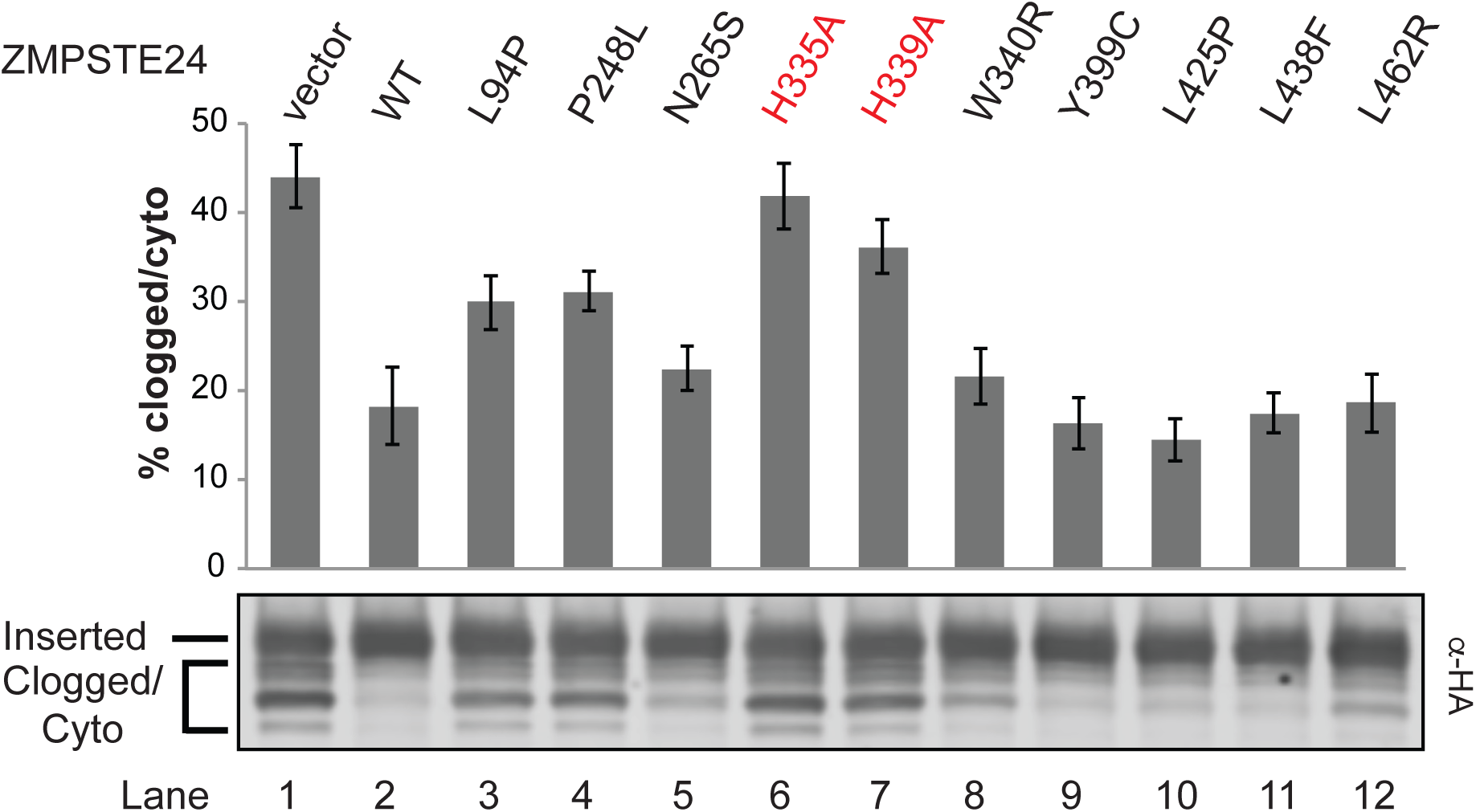
Comparison of ZMPSTE24 mutants for clearance of the “clogger” protein. Strain SM6117 (*ste24*Δ *P_GAL_-Clogger-HA*) transformed with the indicated ZMPSTE24 plasmids was induced to express the clogger protein by addition of galactose, as described in Materials and Methods. Lysates were resolved by SDS-PAGE and probed with α-HA antibodies to detect the clogger. The inserted and clogged or cytoplasmic species are indicated, with the percentage clogged/cytoplasmic graphed on the y-axis. Data shown is the mean ± s.e.m. for five individual experiments. p<0.05 for vector, P248L, H335A and H339A compared to WT.

## DISCUSSION

### A “humanized yeast system” for analysis of ZMPSTE24 cleavage of prelamin A

Gaining an understanding of how the structure of a protein dictates its function is facilitated by assaying the impact of specific mutations on activity, protein stability, and interactions *in vivo*. For proteins involved in disease, this information can also provide invaluable insights into personalized medicine strategies, as exemplified by the customized therapies recently developed to treat cystic fibrosis patients with different disease alleles of the *CFTR* gene (Amaral, 2015). Here we report the development of an *in vivo* humanized yeast system to determine the impact of *ZMPSTE24* disease mutations on the cleavage of prelamin A (Fig. 1, Step 4), the step that is defective in progeroid diseases. Importantly, this yeast system allows us to measure both ZMPSTE24 prelamin A cleavage activity and *in vivo* protein stability, and has the potential to ultimately be scaled up for high throughput analysis of a large number of mutant alleles. Our humanized yeast system retains all of the known requirements observed in mammalian cells (Figs. 2 and 3) in that the prelamin A substrate must be farnesylated and carboxyl methylated for efficient cleavage, and mutation of a residue adjacent to the cleavage site in prelamin A (L647R) abolished processing in yeast, as it does in mammalian cells. Furthermore, mutation of the ZMPSTE24 catalytic motif HEHHX completely blocks prelamin A processing.

We previously used yeast to gain insight into *ZMPSTE24* disease alleles, based on the ability of human ZMPSTE24 to mediate the post-translational maturation of a non-native substrate, the yeast mating pheromone a-factor (Barrowman et al., 2012b). In that study, yeast mating efficiency was measured as a proxy for directly assessing substrate cleavage. We showed that RD null alleles were completely devoid of mating, whereas the five ZMPSTE24 MAD-B missense mutations tested all showed some residual mating activity. That study supported the notion that even a small amount of ZMPSTE24 function diminishes disease severity and is beneficial for patients. However it was not possible to determine whether –AAXing (Fig. 1 Step 2), the final cleavage step (Fig. 1, Step 4), or both were affected, since the *rce1*Δ*ste24*Δ strain used in that study required ZMPSTE24 for both of the a-factor processing events. In the current study, we have developed an improved and completely “humanized yeast” system (both substrate end enzyme are encoded by the human genes). Because Rce1 is present to perform the –AAXing step, the final prelamin A cleavage step is the reaction being measured. Our optimized system not only assays ZMPSTE24’s ability to cleave its *bona fide* substrate human prelamin A, but also provides a good model for what occurs in RD and MAD-B patient cells where ZMPSTE24 is mutated and RCE1 is present.

### ZMPSTE24 missense disease alleles show reduced prelamin A cleavage *in vivo* and define three mutant classes

We tested the eight currently known *ZMPSTE24* missense alleles implicated in progeroid diseases (Table 1), along with two catalytic dead alleles that alter the HEXXH domain (H335A and H339A). While all of the disease mutants exhibit decreased overall prelamin A cleavage compared to wild-type *ZMPSTE24*, all show residual prelamin A cleavage activity, significantly greater than the catalytic dead alleles (Fig. 4 and Table 2). However the mutants vary greatly in the extent of remaining activity (from 6%-57% that of WT). Importantly, four of the ZMPSTE24 mutations (L94P, P248L, W340R and L462R) showed marked decreases in ZMPSTE24 protein levels (14% − 40% of the WT level), suggesting they cause misfolding and subsequent degradation, while for the others, protein stability is only minimally affected. Taking into account both their activity and stability we can divide ZMPSTE24 disease alleles into 3 classes (Table 2): Class I mutations affect mainly cleavage activity (N265S and Y399C), Class II affect mainly protein stability (P248L, W340R), and Class III mutants affect both (L94P, L425P, L438F, and L462R). None of the mutant alleles we tested influence the ER/perinuclear membrane localization of ZMPSTE24-GFP (Fig. S3), or HA-ZMPSTE24 (data not shown).

For the highly unstable mutant proteins (L94P, P248L, W340R and L462R), the UPS is largely responsible for their degradation, since their ZMPSTE24 protein levels are significantly restored when cells are treated with the proteasome inhibitor bortezomib, or when expressed in a *doa10*Δ strain (Figs. 5A and 5C). Similarly, steady-state levels of the other ZMPSTE24 disease alleles, including the least unstable variants (N265S, Y399C, L425P and L438F), increased in the *doa10*Δ mutant, although none as dramatically as P248L (Fig. S4). Quite notably, for the P248L and W340R (Class II) mutants, but not for the L94P and L462R (Class III) mutants, prelamin A cleavage is dramatically restored to near WT levels in the *doa10*Δ strain or following proteasome inhibition using bortezomib (Figs. 5B and 5D). This finding confirms the conclusion that the enzyme function of these two Class II mutant proteins remains largely intact, as also indicated by the “adjusted ZMPSTE24 activity” column in Table 2, despite the presence of mutations that target them for degradation. Indeed, preliminary experiments using purified ZMPSTE24 proteins show that the P248L and W340R variants retain significant prelamin A cleavage activity *in vitro* (EP Carpenter and L. Nie, unpublished observations). P248L was also suggested to retain partial activity in a previous study based on a-factor production (Miyoshi et al., 2008).

It is not uncommon for a mutation that impairs folding only slightly to result in avid protein clearance by the UPS, but for that protein to retain residual function if its degradation is inhibited (Gardner et al., 2005). Thus the protein quality control system is sometimes more aggressive than it needs to be and offers the potential for proteasome inhibitor drugs to be used to ameliorate disease. In the future, it will be of great interest to determine whether ZMPSTE24 protein levels and prelamin A processing can be restored in P248L and W340R disease patient cells by genetic or chemical manipulations of the UPS. Such a finding could point the way to personalized medicine approaches that would differ between patients with Class II mutations and patients with Class I and III mutations that impact protein function. Similar personalized medicine approaches have been successful for cystic fibrosis, in which different drug treatments have been developed for patients with different classes of CFTR mutations (De Boeck and Amaral, 2016). Although not yet directly tested in clinical trials, all patients with mutated versions of ZMPSTE24 are predicted to benefit from farnesyltransferase inhibitors (FTIs), which should render full-length prelamin A unmodified and thus less harmful, as it does for progerin in HGPS (Gordon et al., 2014). However patients with Class II *ZMPSTE24* mutations could potentially improve even more by a combination of FTIs and UPS inhibitors.

### Analysis of genetic dominance for L462R and L438F ZMPSTE24 alleles

Genetic pedigree analysis as well as patient genotypes indicates that generally *ZMPSTE24* mutations are recessive. Thus, disease is usually not manifest in individuals if one of their two *ZMPSTE24* alleles is WT. However two patient mutations, L462R and L438F, have been suggested to be exceptions and could be dominant, since mutations in the second *ZMPSTE24* allele were not identified in these patients by sequence analysis of exons (Table 1). These patients have RD and metabolic syndrome, respectively. Surprisingly however, the healthy mother of the L462R RD patient shared the same apparent genotype *(ZMPSTE24^+/L462^)* as her affected offspring, arguing against dominance (Thill et al., 2008). We propose that L462R is actually recessive and that an as-yet-undetected mutation either inside (and missed, as has previously happened (Navarro et al., 2005)) or outside the coding region (e.g. promoter, intron, etc.) may inactivate the second seemingly WT *ZMPSTE24* allele of the child with RD. A similar explanation may account for disease in L438F patients as well. Supporting this hypothesis and arguing against dominance, we found that an additional wild-type copy of *ZMPSTE24* could efficiently suppress the prelamin A cleavage defect observed in all *ZMPSTE24* disease alleles, including L462R as well as L438F (Fig. S4).

In light of the possibility that both L462R and L438F are actually recessive, an additional aspect of these mutations deserves mention. Most *ZMPSTE24* missense mutations cause MAD-B. However, It is notable that we found that L462R is the most severe of the alleles studied here, while L438F is the least severe, showing significant residual activity (Fig. 4 and Table 2), which may explain why the former leads to the disease RD which is more severe than MAD-B, while the latter leads to metabolic disorder or NAFLD, diseases that are milder than MAD-B (Brady et al., 2017; Dutour et al., 2011; Galant et al., 2016).

### Some, but not all, disease alleles affect ZMPSTE24’s ability to clear clogged translocons

Our studies here suggest that the recently reported role for ZMPSTE24 in clearance of clogged translocons relies on certain features of the protease that may be separable from those required for prelamin A cleavage. As shown previously (Ast et al., 2016), wild-type ZMPSTE24, but not catalytic-dead versions, can efficiently replace yeast Ste24 when challenged with a clogging-prone substrate (Fig. 6). Interestingly, we also show here that some mutants, including Y399C and L462R, show poor prelamin A processing (~25% and 6% of WT, respectively), yet are largely proficient in the unclogging process. It is perhaps not surprising that the two different types of substrates (prelamin A and clogged proteins) might be handled differently. For instance, certain ZMPSTE24 mutations may prevent putative interactions between the protease and the translocon machinery, or in their ability to permit access of the two different types of substrates to the active site. We speculate that like the Sec61 translocon itself, which is known to open laterally to release transmembrane spans (Gogala et al., 2014; Li et al., 2016; Pfeffer et al., 2015), ZMPSTE24 may also have the capacity to do so to facilitate transfer of a clogged substrate from the translocon pore to the ZMPSTE24 protease catalytic site. Isolating mutants specific to each function could help reveal important aspects about these processes. Thus far, we have not identified ZMPSTE24 mutants that are specifically deficient for de-clogging, but such mutants could be sought using our system and will provide further evidence that these activities are separable.

### Utility of our humanized yeast system for structure-function analysis of ZMPSTE24

ZMPSTE24 is an intriguing molecule in terms of basic membrane protein biology, in addition to its importance for understanding progeroid diseases and physiological aging. While ZMPSTE24 has a canonical HEXXH zinc binding motif that can coordinate zinc and mediates catalysis, the ZMPST24 structure is fundamentally different from other proteases, because catalysis occurs within an unusual enclosed intramembrane chamber (Clark et al., 2017; Pryor et al., 2013; Quigley et al., 2013). This novel structure elicits a number of questions concerning ZMPSTE24 enzyme mechanism, prelamin A access and positioning, and why prelamin A is the sole specific substrate known for ZMPSTE24. As a starting point, we focused here on the eight known ZMPSTE24 missense disease alleles. Six of the eight ZMPSTE24 disease alleles lie in residues highly conserved in ZMPSTE24/Ste24 among diverse species (the exceptions are L94 and Y399) and they cluster in two regions of the ZMPSTE24 structure (Fig. 7). Most are at the top of the chamber, near the HEXXH catalytic motif, which coordinates the zinc ion (yellow). It has been suggested that these mutations could affect catalysis directly (N265S), impede substrate binding (L438F and L462R), or substrate entry into the chamber (P248L and W430R) (Quigley et al., 2013). Notably two mutations L94P and Y399C map to the bottom side of the chamber, suggesting a functionally important activity in that region of ZMPSTE24, possibly in farnesyl binding for proper substrate positioning. As discussed above, an important finding here is that some of disease mutations mainly produce an effect by destabilizing the protein, rather than affecting its enzymatic function *per se*.

**Figure 7.**
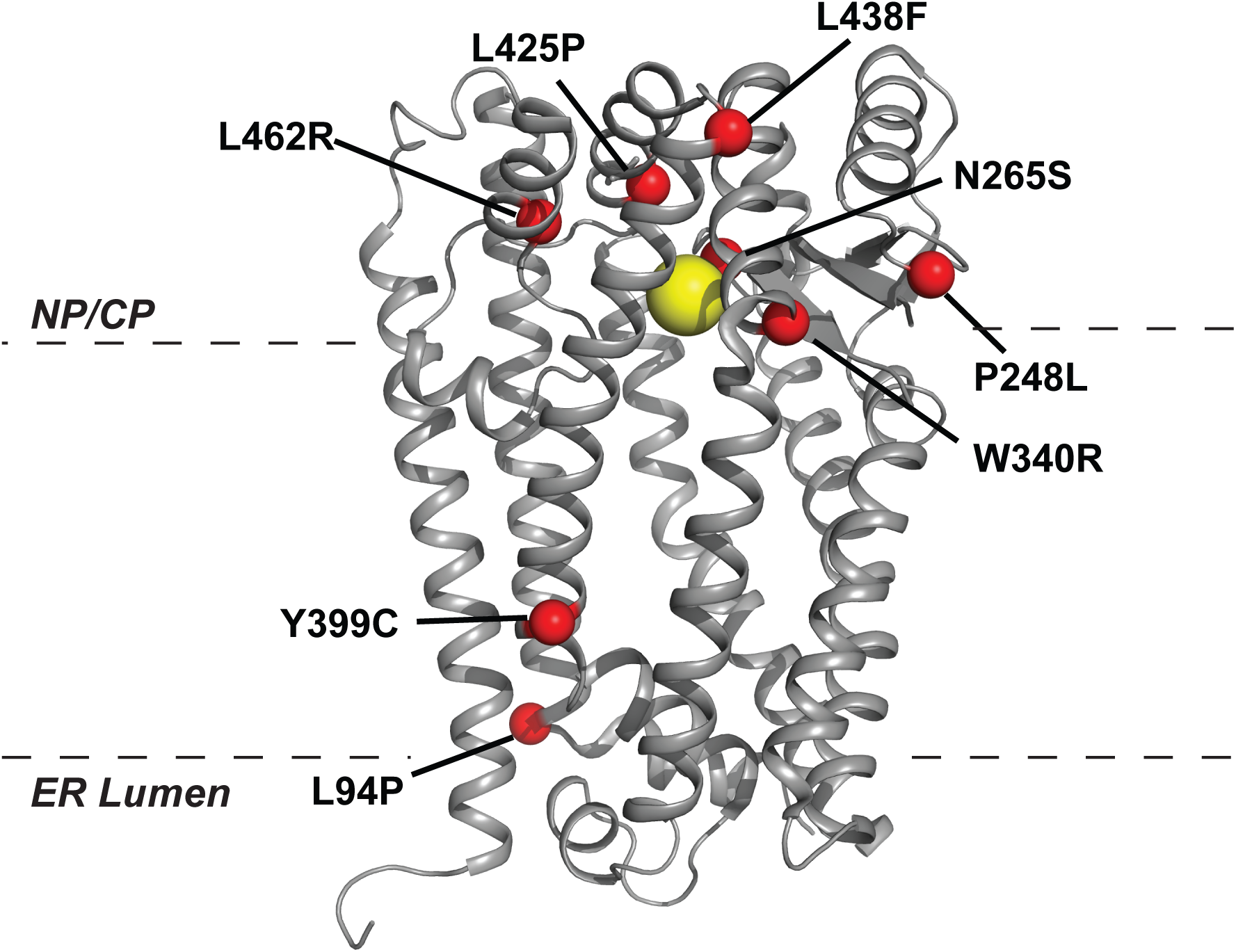
Location of missense disease alleles in the ZMPSTE24 structure. Positions of missense disease alleles in Table 1 are indicated on a ribbon diagram of the ZMPSTE24 structure (PDB entry 2YPT (Quigley et al., 2013)). Yellow ball is zinc at the catalytic site. Dashed lines indicate the approximate delineation of the lipid bilayer, with the ER lumen and nucleoplasm/cytoplasm (NP/CP) indicated.

In the long term, we expect that the humanized yeast system reported here, along with high throughput mutagenesis, will allow us to answer mechanistic questions about how ZMPSTE24 functions and which features of misfolded versions of ZMPSTE24 recruit the UPS-dependent protein quality control machinery, an issue not well understood for any multispanning membrane protein. Our humanized yeast assay will also facilitate dissection of the prelamin A substrate to define a ZMPSTE24 consensus cleavage sequence and to probe the role of farnesyl for ZMPSTE24-mediated cleavage. Ultimately the capacity to perform deep mutational scanning followed by specific screens and selections in yeast (Fowler et al., 2014; Starita and Fields, 2015) will facilitate isolation of separation-of-function alleles, in which ZMPSTE24 residues specific for prelamin A processing, declogging, and antiviral activity can be identified.

## MATERIALS AND METHODS

### Plasmids and strains used in this study

Plasmids used in this study are listed in Table 3. All plasmids were constructed using standard molecular biology techniques, including NEBuilder HiFi Assembly (New England Biolabs) and Quickchange mutagenesis (Stratagene). When mutating ZMPSTE24-containing plasmids, *E. coli* competent cells (Stbl2; Invitrogen) were transformed and grown at 30ΰC. ZMPSTE24 plasmids are *CEN/URA3* containing N-terminally-*His*_10_*HA*_3_-tagged human *ZMPSTE24* expressed from the *PGK1* promoter. For ZMPSTE24-GFP plasmids, a PCR product encoding *GFP* was inserted in between codons 469 and 470 of the *ZMPSTE24* ORF, just subterminal to the C-terminal ER retrieval signal (Barrowman et al., 2008). Plasmid pSM3094 was constructed by subcloning a *SacII-XhoI* fragment containing N-terminally-His_10_HA_3_-tagged yeast *STE24* from pSM1282 (Tam et al., 2001) into the same sites of pRS316. Plasmid pSM3283 is *CEN/HIS3* containing a single Flag-epitope at the N-terminus of ZMPSTE24 expressed from the *PGK1* promoter. Plasmid pSM3204 (expressing mCherry-Scs2™) was constructed by NEBuilder HiFi Assembly of a PCR product from Kp173 (a generous gift of Rong Li, JHU School of Medicine) into pRS315 *(CEN/LEU2)*.

**Table 3.** Plasmids used in this study

| **Plasmid** | **Description** | **Reference** |
| --- | --- | --- |
| pSM171 | pRS313 (CEN, *HIS3*) | (Sikorski and Hieter, 1989) |
| pSM174 | pRS316 (CEN, *URA3*) | (Sikorski and Hieter, 1989) |
| pSM2671 | pRS316::P<sub>PGK1</sub>-10His-3HA-zmpste24N265A | (Barrowman et al., 2012b) |
| pSM2672 | pRS316::P<sub>PGK1</sub>-10His-3HA-zmpste24W340R | (Barrowman et al., 2012b) |
| pSM2673 | pRS316::P<sub>PGK1</sub>-10His-3HA-zmpste24H335A | (Barrowman et al., 2012b) |
| pSM2676 | pRS316::P<sub>PGK1</sub>-10His-3HA-zmpste24P248L | (Barrowman et al., 2012b) |
| pSM2677 | pRS316::P<sub>PGK1</sub>-10His-3HA-ZMPSTE24 | (Barrowman et al., 2012b) |
| pSM2982 | pRS316::P<sub>PGK1</sub>-10His-3HA-zmpste24L94P | (Barrowman et al., 2012b) |
| pSM2984 | pRS316::P<sub>PGK1</sub>-10His-3HA-zmpste24L438F | (Barrowman et al., 2012b) |
| pSM3094 | pRS316::P<sub>PGK1</sub>-10His-3HA-STE24 | This study |
| pSM3162 | pRS316::P<sub>PGK1</sub>-10His-3HA-zmpste24H339A | This study |
| pSM3185 | pRS316::P<sub>PGK1</sub>-10His-3HA-zmpste24L425P | This study |
| pSM3186 | pRS316::P<sub>PGK1</sub>-10His-3HA-zmpste24Y399C | This study |
| pSM3204 | pRS315::P<sub>TDH3</sub>-mCherry-SCS2-Tm | This study |
| pSM3283 | pRS313::P<sub>PGK1</sub>-Flag-ZMPSTE24 | This study |
| pSM3317 | pRS316::P<sub>PGK1</sub>-10His-3HA-zmpste24L462R | This study |
| pSM3173 | YIP-TRP1::NatMX-P<sub>PRC1</sub>-10His-3myc-LMNA(431-664) | This study |
| pSM3177 | YIP-TRP1::NatMX-P<sub>PRC1</sub>-10His-3myc-LMNA(431-664, L647R) | This study |
| pSM3178 | YIP-TRP1::NatMX-P<sub>PRC1</sub>-10His-3myc-LMNA(431-646) | This study |
| pSM3360 | YIP-TRP1::NatMX-P<sub>PRC1</sub>-10His-3myc-LMNA(431-664, C661S) | This study |
| pSM3429 | pRS316::P<sub>PGK1</sub>-10His-3HA-ZMPSTE24-GFP469 | This study |
| pSM3430 | pRS316::P<sub>PGK1</sub>-10His-3HA-zmpste24L94P-GFP469 | This study |
| pSM3431 | pRS316::P<sub>PGK1</sub>-10His-3HA-zmpste24P248L-GFP469 | This study |
| pSM3455 | pRS316::P<sub>PGK1</sub>-10His-3HA-zmpste24H335A-GFP469 | This study |
| pSM3456 | pRS316::P<sub>PGK1</sub>-10His-3HA-zmpste24Y399C-GFP469 | This study |
| pSM3457 | pRS316::P<sub>PGK1</sub>-10His-3HA-zmpste24L462R-GFP469 | This study |

Plasmid pSM3173 is an integrating vector derived from Kp173. Briefly, PCR-generated fragments from the *PRC1* promoter (−800 to −1), *His_10_-myc_3_* and human *LMNA* (corresponding to amino acids 431-664) were recombined *in vitro* with a PCR-generated gapped Kp173 using NEB HiFi Assembly. Plasmids pSM3177 (L647R) and pSM3360 (C661S) were generated with mutagenic oligos and Quickchange mutagenesis using pSM3173 as template. Plasmid pSM3178 that expresses mature lamin A was constructed by placing a stop codon after amino acid Y646 using NEB HiFi Assembly. All manipulations with *LMNA* sequences used DH5α or NEB5 cells (NEB) for propagation. All integrating vectors in this study recombine at the *TRP1* locus by selecting with nourseothricin (Nat). Plasmid sequences and maps available upon request.

Yeast strains used are listed in Table 4. To integrate *LMNA* constructs, integrating plasmids were linearized by EcoRV digestion and transformed into *ste24*Δ (SM4826) cells by standard yeast lithium acetate protocols. Transformants were selected on YPD containing 100 µg/ml nourseothricin. To generate the double mutants *ste24*Δ*doa10*Δ and *ste24*Δ*ste14*Δ, diploids were made by crossing single mutant strains of opposite mating types and the double mutants identified following sporulation and tetrad dissection. Strain SM6117 (*ste24*Δ *P_GAL1_-PDI1-DHFR-Nglyc-3HA* (clogger) was used previously (Ast et al., 2016). ZMPSTE24-expressing plasmids were transformed into strains and selected on minimal SC-Ura or SC-Ura-His plates.

**Table 4.** Yeast strains used in this study

| **Strain** | **Genotype** | **Reference** |
| --- | --- | --- |
| SM4826 | *ste24::KanMX met15Δ0 his3Δ1 leu2Δ0 ura3Δ0 Mata* | Deletion collection |
| SM6117 | *ste24::Hyg<sup>r</sup> HO::P<sub>GAL1</sub>-PDI1-clogger-3HA met15Δ0 his3Δ1 leu2Δ0 ura3Δ0 Mata* | (Ast et al., 2016) |
| SM6158 | *ste24::KanMX met15Δ0 his3Δ1 leu2Δ0 ura3Δ0 Mata TRP1::NatMX-P<sub>PRC1</sub>-10His-3myc-LMNA(431-664) Mata* | This study |
| SM6159 | *ste24::Hyg<sup>r</sup> pdr5::KanMX met15Δ0 his3Δ1 leu2Δ0 ura3Δ0 TRP1::NatMX-P<sub>PRC1</sub>-10His-3myc-LMNA(431-664) Mata* | This study |
| SM6173 | *ste24::KanMX met15Δ0 his3Δ1 leu2Δ0 ura3Δ0 Mata TRP1::NatMX-P<sub>PRC1</sub>-10His-3myc-LMNA(431-664, C661S) Mata* | This study |
| SM6177 | *ste24::KanMX met15Δ0 his3Δ1 leu2Δ0 ura3Δ0 Mata TRP1::NatMX-P<sub>PRC1</sub>-10His-3myc-LMNA(431-664, L647R) Mata* | This study |
| SM6178 | *ste24::KanMX met15Δ0 his3Δ1 leu2Δ0 ura3Δ0 Mata TRP1::NatMX-P<sub>PRC1</sub>-10His-3myc-LMNA(431-646) Mata* | This study |
| SM6184 | *ste24::KanMX doa10::KanMX met15Δ0 his3Δ1 leu2Δ0 ura3Δ0 Mata TRP1::NatMX-P<sub>PRC1</sub>-10His-3myc-LMNA(431-664) Mata* | This study |
| SM6187 | *ste24::KanMX ste14::KanMX lys2Δ0 his3Δ1 leu2Δ0 ura3Δ0 Mata TRP1::NatMX-P<sub>PRC1</sub>-10His-3myc-LMNA(431-664) Matα* | This study |

### Yeast prelamin A cleavage assay

Typically, strains grown overnight in minimal medium (0.67% yeast nitrogen base, 0.5% ammonium sulfate, 2% glucose, supplemented with appropriate amino acids and supplements) were back diluted in fresh medium for 4-6 hours. Cells (1.5-2 O.D._600_ cell equivalents) were pelleted, washed in water and lysed using NaOH pre-treatment and SDS protein sample buffer (Kushnirov, 2000) at 65ΰC for 10-15 min. For analysis, lysates were centrifuged at 21k x g for 2 min, and the supernatant (0.3 OD cell equivalents per lane) were resolved on 10% SDS-polyacrylamide gels. Proteins were transferred to nitrocellulose (Bio-Rad Transblot Turbo), and the membrane was blocked using Western Blocking Reagent (Roche). Lamin proteins were detected using mouse anti-myc antibodies (clone 4A6, Millipore cat # 05-724; 1:10,000 dilution) decorated with goat anti-mouse secondary IRDye 680RD antibodies (LI-COR). Blots were re-probed using rat anti-HA (clone 3F10, Roche cat # 11867423001; 1:10,000 dilution) to detect ZMPSTE24 and rabbit anti-Sec61 (1:10,000 dilution) as a loading control (a generous gift of Dr. Randy Schekman, UC, Berkeley), and visualized using goat anti-rat IRDye 680RD and goat anti-rabbit IRDye 800CW secondary antibodies (LI-COR). Prelamin A cleavage was calculated using ImageStudio Lite (LI-COR) by quantifying mature lamin A signal compared to total lamin A signal (prelamin A + mature lamin A). ZMPSTE24 protein levels were quantified by measuring the HA signal in the entire region that contains both ZMPSTE24 bands and the intervening smear and normalizing this signal to the Sec61 loading control signal. Statistical analyses were performed by unpaired, two-tailed t-test (using Microsoft Excel), with p<0.05 considered to be significant.

### Clogger assay

Translocon clogging was examined essentially as previously described (Ast et al., 2016). Strain SM6117 transformed with vector or ZMPSTE24-expressing plasmids were grown overnight in SC-Ura with 2% sucrose as the carbon source. Strains were back diluted in the same medium for 3 hours, and then induced by adding galactose to 2.5% for six hours prior to collecting cells. SDS-PAGE, western transfer and blocking were done as described for prelamin A cleavage. Clogger protein was detected using rat anti-HA (3F10, Roche) and LI-COR secondary antibodies. Inserted, and clogged/cytoplasmic forms were quantified using ImageStudio Lite (LI-COR).

### Proteasome inhibition

To test the effect of proteasome inhibition on ZMPSTE24 protein levels and prelamin A cleavage, strain SM6159 (*ste24*Δ*pdr5*Δ *P_PRC1_-10His-3myc-LMNA_CT_*) transformed with the indicated ZMPSTE24 alleles were grown to log phase in SC-Ura medium, and treated with either DMSO or 20 µM bortezomib (from 30mM stock in DMSO, a generous gift from Peter Espenshade, JHU School of Medicine) for 4 hrs at 30ΰC prior to lysate preparation, SDS-PAGE and western blotting as described earlier. MG-132 was also tested with similar results (data not shown). The *pdr5*Δ mutation was introduced to enhance the efficacy of drug treatment as previously described (Collins et al., 2010; Sung et al., 2016).

## ACKNOWLEDGMENTS

This work was funded by grants R01 GM041223 (to SM) and R01 GM106082 (to CAH) from the National Institutes of Health. EPC and LN are funded for this work by UK Medical Research Council grant number MR/L017458/1. EPC is also funded by the Structural Genomics Consortium (SGC), which is a registered charity (number 1097737) that receives funds from AbbVie, Bayer Pharma AG, Boehringer Ingelheim, the Canada Foundation for Innovation, Genome Canada, GlaxoSmithKline, Janssen, Lilly Canada, Merck & Co., the Novartis Research Foundation, the Ontario Ministry of Economic Development and Innovation, Pfizer, São Paulo Research Foundation-FAPESP, Takeda, EU/EFPIA Innovative Medicines Initiative (IMI) Joint Undertaking (ULTRA-DD grant n° 115766) and the Wellcome Trust [092809/Z/10/Z].

---

**Figure S1.**
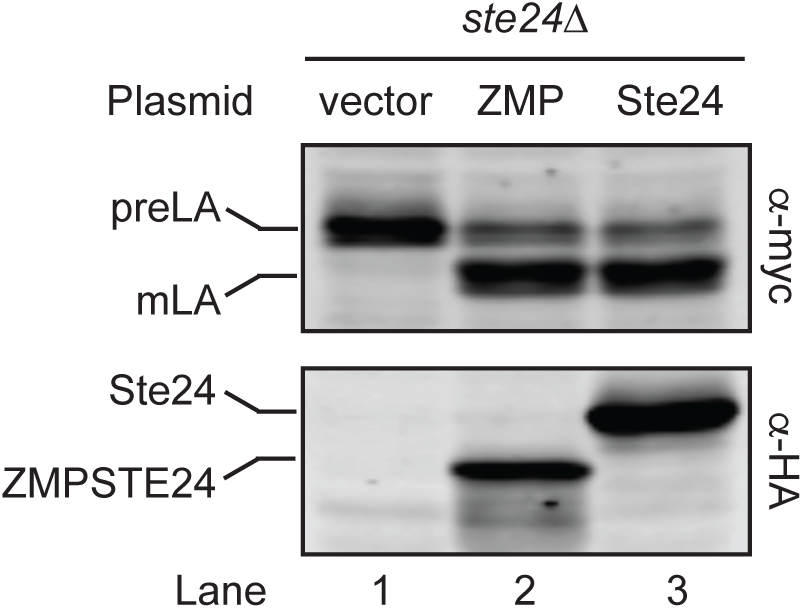
Yeast Ste24 can catalyze prelamin A processing as efficiently as human ZMPSTE24. Lysates were prepared from SM6158 (*ste24*Δ *myc-LMNA_CT_*) cells transformed with vector only (Lane 1), pSM2677 (human *ZMPSTE24*, Lane 2) or pSM3094 (yeast *STE24*, Lane 3) and subjected to SDS-PAGE and Western blotting with α-myc antibodies to detect processing of prelamin A and α-HA antibodies to detect the ZMPSTE24 or Ste24 proteins, which both contain an N-terminal 10His-3HA epitope tag and are expressed from the same promoter. Although similar in size, ZMPSTE24 (475 amino acids) and Ste24 (453 amino acids) migrate significantly different by SDS-PAGE, with Ste24 surprisingly migrating more slowly than ZMPSTE24, despite its smaller size. However, anomalous SDS-PAGE migration patterns are not uncommon among membrane proteins, due to unpredictable differences in their detergent binding properties (Newman et al., 1981; Rath et al., 2009).

**Figure S2.**
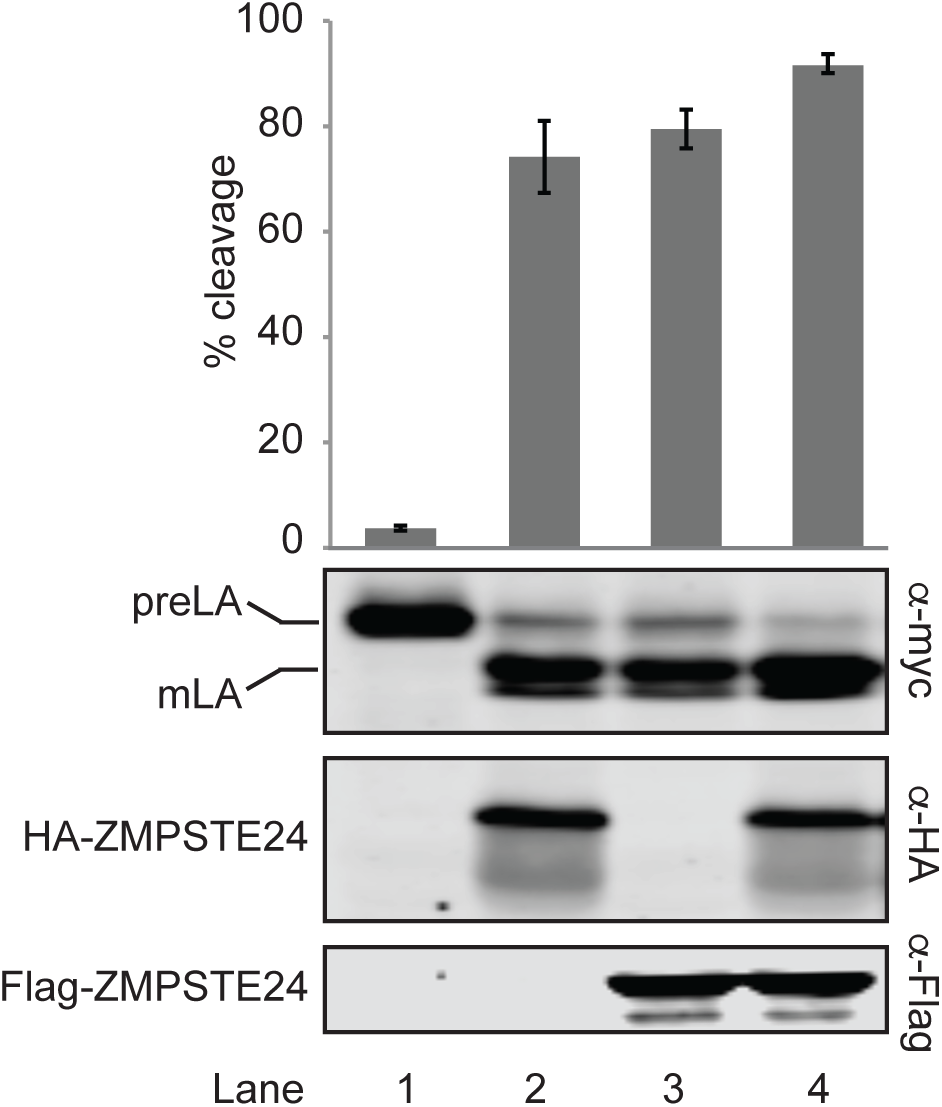
The extent of prelamin A processing depends on the amount of ZMPSTE24 cells express. Lysates were prepared from strain SM6158 (*ste24*Δ *myc-LMNA_CT_*) containing empty vectors only (pRS313 and pRS316; Lane 1), HA-ZMPSTE24 and empty vector (pSM2677 and pRS313; Lane 2), Flag-ZMPSTE24 and empty vector (pSM3283 and pRS316; Lane 3) or HA-ZMPSTE24 and Flag-ZMPSTE24 (pSM2677 and pSM3283; Lane 4) and were analyzed by western blotting as described. Prelamin A cleavage is expressed as the mean ± s.d. from three independent trials. p<0.05 for vectors and one copy compared to two copies of ZMPSTE24. With a single copy of ZMPSTE24 expressed from the *PGK1* promoter, the prelamin A model substrate undergoes 65-80% processing (lanes 2 and 3), which increased to 90% with two copies (lane 4). It is possible that 100% processing cannot be achieved because a small percentage of prelamin A is not farnesylated.

**Figure S3.**
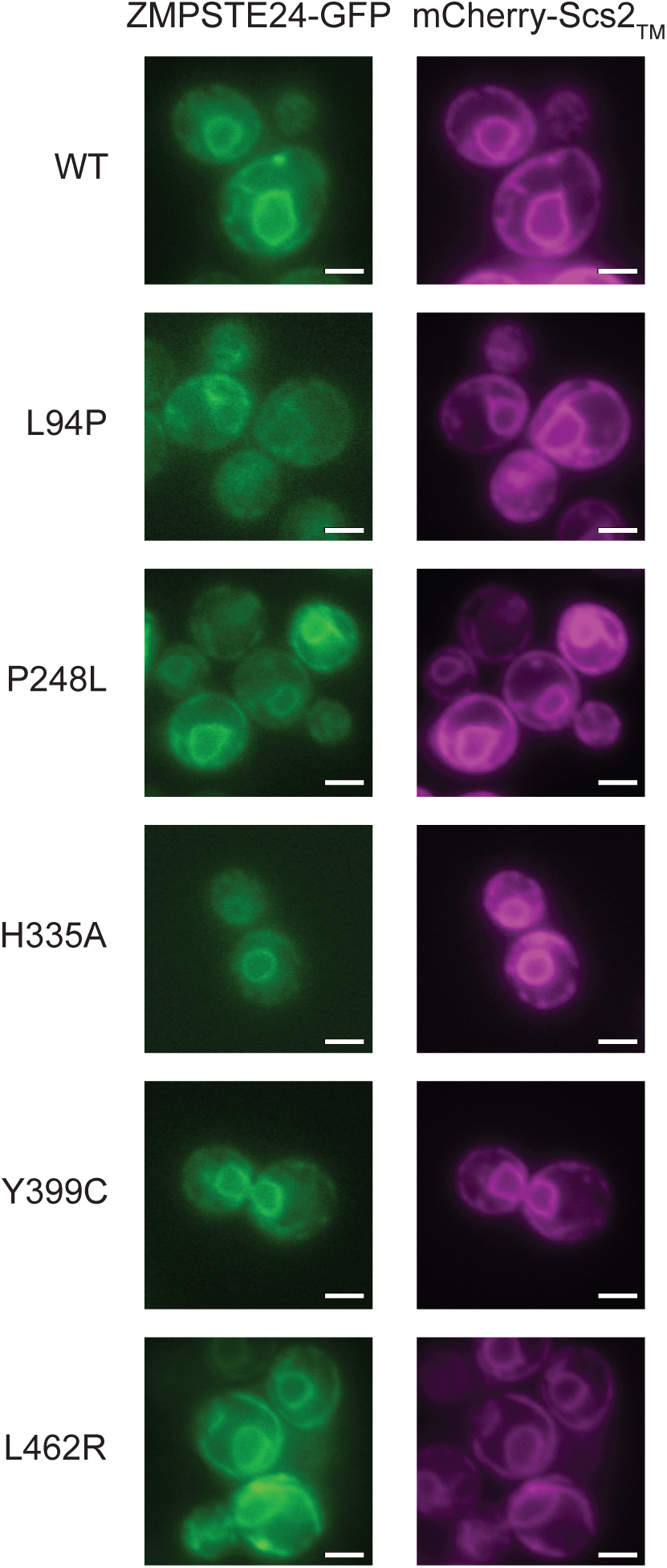
WT and ZMPSTE24-GFP variants localize to the perinuclear and cortical ER membranes. Strain SM4826 (*ste24*Δ) was transformed with mCherry-Scs2TM (pSM3204) and the indicated ZMPSTE24-GFP variants were visualized by fluorescence microscopy. mCherry-Scs2TM is an ER marker. Shown are the ZMPSTE24 mutants having the most severe prelamin defects. All HA-ZMPSTE24 mutants show no obvious localization changes by indirect immunofluorescence (data not shown). Scale bar, 2μM.

**Figure S4.**
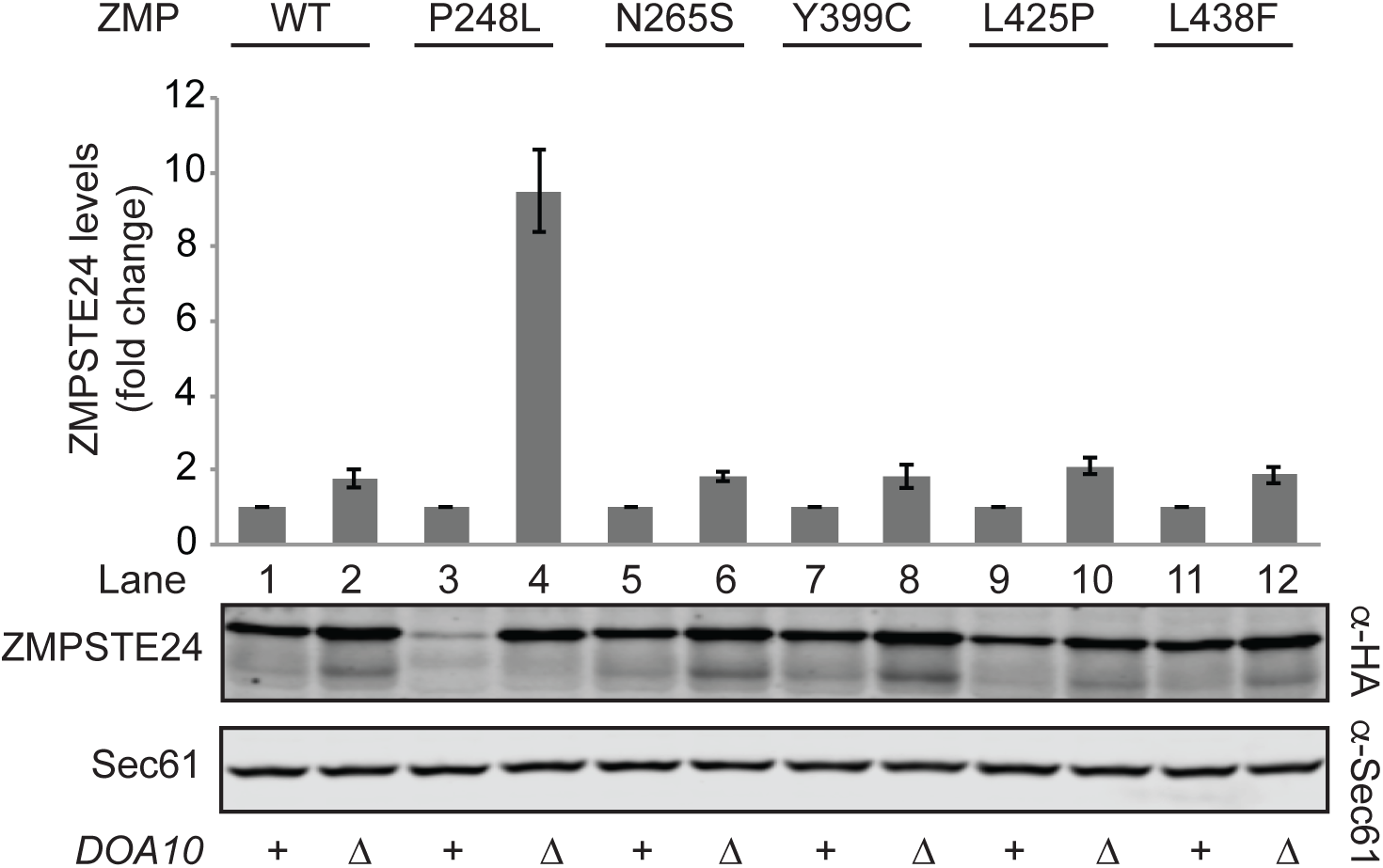
ZMPSTE24 protein levels are increased in the *doa10*Δ mutant. Strains SM6158 (*ste24*Δ *myc-LMNA_CT_*) or SM6184 (*ste24*Δ*doa10*Δ *myc-LMNA_CT_*) expressing the indicated ZMPSTE24 variants were analyzed by SDS-PAGE and western blotting as described in Figure 5. ZMPSTE24 steady-state levels were normalized to the loading control Sec61, and the mean ± s.d. for three independent experiments is shown. p<0.05 for all ZMPSTE24 variants between WT and *doa10*Δ cells.

**Figure S5.**
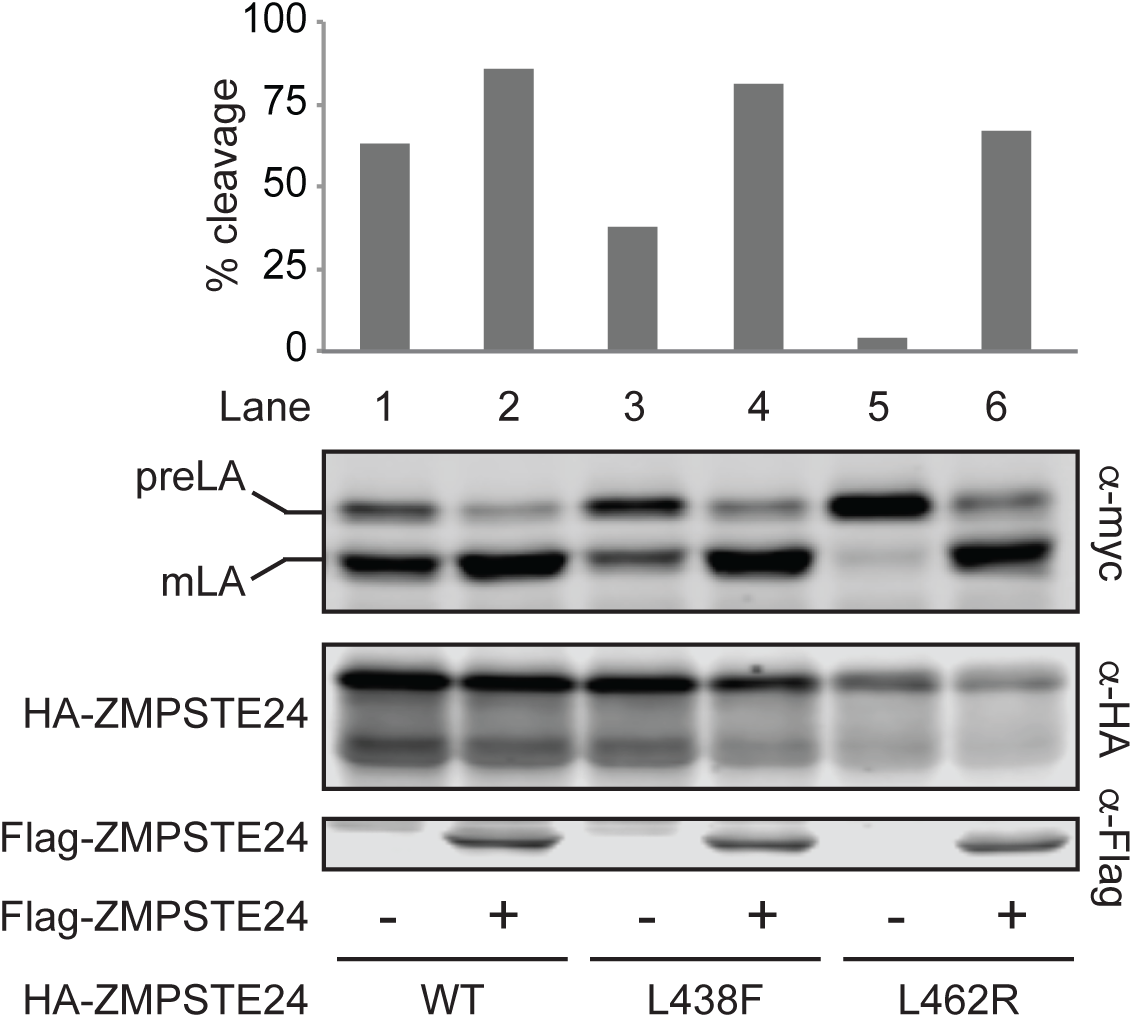
ZMPSTE24 disease alleles do not exhibit dominance in the yeast prelamin A cleavage assay. Strain SM6158 (*ste24*Δ *myc-LMNA_CT_*) transformed with the indicated HA-ZMPSTE24 disease alleles and either pRS313 (vector only; −) or pSM3283 (which expresses WT Flag-ZMPSTE24; +) were assayed for in vivo prelamin A cleavage. Lysates were resolved by SDS-PAGE and proteins detected by immunoblotting against myc (prelamin A), HA (HA-ZMPSTE24) and Flag (Flag-ZMPSTE24). So long as at least one ZMPSTE24 allele is WT (lanes 2, 4 and 6) processing of prelamin A is close to 70%, even if a mutant allele is present.

